# Signatures of Brain Network Alteration in Psychogenic Non-Epileptic Seizures: A Rest-EEG Study Based on Power Spectral Density and Phase Lag Index

**DOI:** 10.1101/2021.10.20.464353

**Authors:** Giuseppe Varone, Wadii Boulila, Michele Lo Giudice, Bilel Benjdira, Nadia Mammone, Cosimo Ieracitano, Kia Dashtipour, Sabrina Neri, Sara Gasparini, Francesco Carlo Morabito, Amir Hussain, Umberto Aguglia

## Abstract

The main challenge in the clinical assessment of Psychogenic Non-Epileptic Seizures (PNES) is the lack of an electroencephalographic marker in the electroencephalography (EEG) readout. Although decades of EEG studies have focused on detecting cortical brain function underlying PNES, the principle of PNES remains poorly understood. To address this problem, electric potentials generated by large populations of neurons were collected during the resting state to be processed after that by Power Spectrum Density (PSD) for possible analysis of PNES signatures. Additionally, the integration of distributed information of regular and synchronized multi-scale communication within and across inter-regional brain areas has been observed using functional connectivity tools like Phase Lag Index (PLI) and graph-derived metrics. A cohort study of 20 PNES and 19 Healthy Control subjects (HC) were enrolled. The major finding is that PNES patients exhibited significant differences in alpha-power spectrum in brain regions related to cognitive operations, attention, working memory, and movement regulation. Noticeably, we observed that there exists an altered oscillatory activity and a widespread inter-regional phase desynchronization. This indicates changes in global efficiency, node betweenness, shortest path length, and small worldness in the delta, theta, alpha, and beta frequency bands. Finally, our findings look into new evidence of the intrinsic organization of functional brain networks that reflects a dysfunctional level of integration of local activity across brain regions, which can provide new insights into the pathophysiological mechanisms of PNES.

## 1 Introduction

Although decades of intensive research, one of the most clinical challenges is related to the management of subjects reporting occurrences of paroxysmal episodes of loss of consciousness or altered awareness. Psychogenic Non-Epileptic Seizures (PNES) are rapid convulsive changes in behavior or consciousness that mimic epilepsy but without electrophysiological bio-markers, which characterize an epileptic seizure in EEG readouts. The investigation of these medical cases, despite extensive and exhaustive clinical inquiries, may remain unfathomable because they largely focused on an insufficient clinical history of the events’ description. Furthermore, the lack of an alternative and robust electrophysiological marker reduces the effectiveness of disentangling the PNES disorder from healthy control subjects in EEG readouts. In well-equipped clinical facilities, the diagnosis of PNES can be strengthened using video-EEG monitoring, with particular mention for those cases where a psychogenic event is captured during spontaneous occurrences or externally triggered episodes [1]. However, video-electroencephalography analysis is expensive, ethically controversial, not commonly available, time and resource-consuming, and finally also prone to bias due to subjective interpretation. In spite of this, clinicians routinely face a demand of differential diagnoses of PNES in an EEG readout free from electrophysiological features. An early and accurate diagnosis of PNES represents an important challenge for patients, proxies, and caregivers. In fact, under diagnosis (or epilepsy misdiagnosis), patients may be subject to a risk of inappropriate and potentially harmful treatment, with a possible impact on patients’ health and placing high burdens on healthcare systems [2]. This makes discrimination of PNES via EEG readout a challenging task for clinicians, especially for cases of false alarms leading to harmful treatments.

In this pasomewhatpropose a methodological pipeline based on Power Spectral Density (PSD) and Phase Lag Index (PLI). The PSD is used to grasp the signature of spectral behaviors, and the PLI, concurrently with graph-driven metrics, is used to grasp the statistical inter-dependencies between two anatomically distinct electrophysiological time-series. The first goal is to investigate the mechanism underlying information transfer among different cortices locations. The repertoire of spontaneous resting-state cortical activity emerges either as a result of the embedded oscillatory activity of neurons and neuronal populations or as an ongoing dynamically changing in the cluster of interconnected neurons [3]. This leads to short-term network connection of boundless neurons. Recent studies on inter-regional brain network communication showed that neural activity is characterized by dynamic fluctuation and synchronization of oscillatory activity between neuronal populations within a millisecond of delay. Neurophysiologically, the intrinsic good hallmarks of brain cooperation rely on the prowess of multiple functionally specialized areas interacting rapidly and effectively. Thus, if two or more cortical generators promote neural oscillations synchronized in frequency and/or in phase, a higher gain of information is transferred; thus a more efficient communication occurs. Hence, an emerging idea is that a healthy brain requires an optimal balance between functional integration and functional differentiation in cortical networks. To this end, studying the rhythmic and phase characteristics of electrophysiological data can reveal the neural infrastructure that underlies both healthy and PNES disordered subjects as potential phenomena of transient inter-regional synchronization. To date, a substantial amount of evidence exists for different brain diseases, but differently from the former, the rest EEG data of PNES have not been extensively investigated so far. In terms of spontaneous scalp EEGs, a general trend is to investigate the redistribution of power in the frequency bands to grasp the peak of frequency or dominant band. Fourier-based power spectral density (PSD), to characterize the frequency distribution of resting-state PNES via EEG data, was used only in [4]. Functional connectivity within particular resting-state networks can be used to study local organization and global integrated and spatially distributed architecture. In the literature, we found a few EEG studies that have focused on derivative tools from graph theory to characterize the synchronization topologies of the neural network of PNES brain systems [5–8]. Noteworthy, we found different techniques applied to study the whole-head neural network synchronization of PNES, including coherence analysis [8], multivariate phase synchronization [5], synchronization likelihood [9] and functional connectivity [10–13], with somewhat contrasting results. According to previous studies, we grasp that there are two major critical issues in the functional brain network of PNES that need to be further investigated.First, despite different studies, there is no gold standard on the binarization approach to use to estimate functional network interaction. Second, there are conflicting results in connectivity analyses of PNES. These differences could be related to the impact of subjects and environment and may be due to the use of different methods, such as electrode density, reference technique, coupling methods and binarization approaches, etc. In addition, we aimed to deal with the following points:

- Investigating possible neural reorganizations including intrinsic alterations in resting-state brain networks oscillations of patients with PNES using PSD analysis.
- Determining whether functional connectivity alterations of PNES subjects could be associated with specific areas which lead to regional network dysfunctions in local oscillations as well as inter-regional synchronization.

The remainder of the paper is structured as follows: first, in Section 2, we review the state-of-the-art in PNES power spectrum and network analysis. Then, in Section 3, we discuss and provide details on the proposed method, including data preprocessing and the techniques used. Next, In Section 5, we present the experimental results on extracting important features from rest-EEG data of PNES subjects. Finally, Section 6 concludes this paper with a breakdown of the proposed method and a discussion of future work towards improving the models.

## 2 Related Works

During the last decades, a wealth of studies have focused on scalp-EEG data, recorded in resting-state condition, to identify criteria that can differentiate PNES disorders based on its phenomenology [14]. Many of these studies provide a window into neural network oscillations [5, 11], functional connectivity [7], loss of integration [15], and dissociation (lost of coherence) [10]. Additionally, in many of these studies is hypothesized that PNES symptoms could be related either to neural network instability, [5–7, 16] due to a mechanism of a time-limited interruption in neural networks connectivity and disconnection between cortical and sub-cortical systems [7, 17], or a time-limited increment in functional connectivity between limbic and motor regions [11, 16].

Knyazeva et al. [5] used multivariate phase synchronization to reconstruct the synchronization of the whole-head topography between cortical regions in PNES. The results show a right prefrontal reduction and alteration in synchronization across all frequency bands with hypersynchronization over the left frontotemporal, parietal-temporal networks, along with hypo synchronization in both right and left frontal regions. Using a graph-theoretical approach, Barzegaran et al. [6] studied the function of cortical networks in patients with PNES using Laplacian-transformed time-series and cross-correlation method, founding that patients with PNES have close to normal local and global connectivity and small-world structure. Xue et al. [8] applied clustering coefficients (local connectivity) and global efficiency to check the coherence of neural networks for patients with PNES. Clustering coefficients and global efficiency were lower in all four frequency bands, but this difference was statistically significant only in the gamma band. In addition, the authors found that PNES had decreased long linkage between the frontal region and posterior brain areas in the gamma band. Barzegaran et al. [7] used multivariate phase synchronization to grapple lagged functional connectivity—between cortical and subcortical regions. Their results observed a decreased functional connectivity between the basal ganglia and limbic, prefrontal, temporal, parietal, and occipital regions, in the alpha band. Umesh et al. [9] performed power spectral and lagged phase synchronization analysis in the gamma band using sLORETA software. The results display a higher power spectrum in the gamma band in the right temporal region. In addition, the authors found a decreased gamma band in the right parietal cortex, posterior cingulate cortex, and superior temporal gyrus. The analysis of connectivity showed a reduced intracortical lagged coherence between the right posterior cingulate gyrus and right middle temporal gyrus. Meppelink et al. [18] found EEG spectral power changes in PNES before seizure events. In addition, the authors found that PNES subjects showed a decrease in beta power (desynchronization) at 5–6 seconds before the PNES event. Arikan et al. [19] used a quantitative EEG (qEEG) and Fast Fourier Transformation (FFT) to investigate the spectral power across all frequency bands in adults with PNES. In this paper, the authors found increased power around the C3 EEG scalp sensor in the beta band, and additionally statistically meaningfully increased gamma power was found in P3 sensor. Amiri et al. [20] applied a graph network analysis for binary functional connectivity by extracting graph-theoretical measures like a nodal degree in cortical and subcortical regions of the brain. Their experimental findings suggest that the functional connectivity can be altered in individuals with PNES. Areas with low connectivity may be involved in emotion processing and movement regulation, whereas areas with higher connectivity may play a role in the inhibition of unwanted movements and cognitive processes.

In the end, recent emerging evidence from neuroimaging studies suggests that neural network instability may be involved in PNES generation and maintenance [16]. Some studies suggest that the neural networks of patients with PNES, when compared to those of healthy controls, maybe less resilient in the face of additional demands and more incline to aberrant shift in functional connectivity—resulting in PNES.

## 3 Materials and Methods

### 3.1 Participants

In this study, we retrospectively reviewed EEG datasets of 80 patients (20 males and 60 females, aged 18–68 years, mean 38.41 years, and SD 16.80), who were hospitalized in the Regional Epilepsy Center, Great Metropolitan Hospital Bianchi Melacrino Morelli, Reggio Calabria, in Italy during the years 2016-2019. All data were collected as part of a routine clinical workup. The protocol of the study was approved by the Medical University of Magna Graecia of Catanzaro (Italy) and all analyses were performed in accordance with the approved guidelines. The collected data have been cleared of identification references related to patients. The subjects with PNES were selected based on the 2013 ILAE Non-Epileptic Seizures Task Force Recommendations [21].

Within the studied group, 60 of the subjects recruited from inpatient or scheduled clinical visits were excluded for use of psychotropic drugs (n = 40), left-handedness (n = 12), or because EEG traces could not be evaluated due to the strong presence of artifacts (n = 8). Finally, 20 EEGs from 20 patients with PNES (7 males, mean 35.05 years, SD 13.07) were included. Three inclusion criteria were considered: 1) at least a single typical PNES episode was recorded by video EEG, and EEG did not show any epileptiform patterns or ictal features; 2) patients had no clinical history of brain disease; 3) patients had no obvious irregularity in structural Magnetic Resonance Imaging (MRI) examinations. In addition, two exclusion criteria were selected: 1) subjects with neurological comorbidities (e.g., epilepsy); 2) subjects with simulated or true psychiatric disorders (e.g., mood and anxiety disorders, schizophrenia, and psychosis).

We also enrolled healthy right-handed controls (HC) without a history of neurological disease, seizures, or sleep disorders who did not consume caffeine and were not taking any medicine until the EEG recording. The control group comprised 19 EEGs from 19 healthy right-hand subjects (5 males, mean age: 35.05 years, SD 13.07). The choice to limit the study to right-handed subjects was due to the uncertain hemispheric dominance in the left-handed brain, which might have influenced results. The Edinburgh Handedness Questionnaire (EHQ) was used to assess hand dominance [22]. We focused on a large cross-sectional community sample of 39 adults in the age range from 18 to 68, to overcome limitations related with age-related changes in spectral EEG content and network density connectivity in a wide frequency band. The sensitivity power analysis conducted on G* Power [23] revealed that our cohort size was large enough to have a statistical power (1 – *β*) of 0.60 to detect significant differences (*α* = 0.05) between subjects with an effect size of 0.65 (Cohen’s d for Welch test).

### 3.2 EEG Recording and Preprocessing

The EEGs were collected in an electromagnetically shielded and light-controlled room where the participants were seated comfortably with closed eyes in a lab chair. The data were recorded according to a standard 20-minute clinical protocol including rest condition using a 19 channels EEG system (Micromed, Italy), having a frequency response from DC to 100 Hz ( which attenuates by 40 dB per decade). The signal was processed through an analog 0.5Hz high-pass filter and digitized with a 12-bit analogical to digital converter (ADC) at 512 Hz. The average duration of recording was 20.2 minutes and has a range of 19.5–22.1 minutes. The recording transducers, Ag/AgCl ring electrodes, were accommodated according to the international 10-20 system with channel layout: Fp1, Fp2, F3, F4, C3, C4, P3, P4, O1, O2, F7, F8, T3, T4, T5, T6, Fz, Cz, and Pz, on G2 (between electrodes Fz and Cz). The electrode-skin impedance was less than 20 kΩ. EEG signals were down-sampled to 256 Hz, digitally band-pass filtered (0.5–200 Hz, 24 dB/octave), and further we applied notch filtered to reduce power line interference. Finally, stored on a local server for further analysis.

The analysis of participants’ EEG data was performed with custom-built scripts in MATLAB (R2021, Mathworks Inc., Natick, MA,USA) and EEGLAB toolbox [24]. First, the collected EEG data were visually inspected to detect bad electrodes or artifacts (e.g., jaw clenching or head-scratching), additionally high-frequency artifacts, such as muscle and high-amplitude slow wave were rejected by means of an automatic algorithm with an initial threshold fixed at 80 *μ*V. Second, the bad channels identified in the previous step were interpolated and a common average referencing was performed. Third, the data were further preprocessed by 0.5 Hz high-pass filter, and FastICA independent component analysis (ICA) was performed to remove components related to eye blinking, eye movement, muscle activity, heartbeat artifacts and instrumental noise. All ICA components were inspected visually and manually selected for rejection (3.41 ± 0.19). Additionally, to validate the selection of ICA components we apply a ICLabel classification [25]. Fourth, twenty-minute of artifacts free scalp EEG data were epoched from 0 to 5 sec into 240 non-overlapping chunks of equal length. Fifth, each individual channel was digitally band-passed, using zero-phase Blackman-windowed FIR filters (cut-off frequencies = **f**_*low*_(f=0.5 Hz) and **f**_*high*_(f=45 Hz), transition bandwidth = 2Hz), to select the EEG rhythm range of interest. Finally, for each epoch we have applied a third-order zero-phase shift Butterworth filter to capture the significant electrophysiologic sub-bands: delta (0.5–3.5 Hz), theta (4–7.5 Hz), alpha (8–13 Hz), and beta (13–32 Hz). The proposed pipeline is graphically presented in Figure. 1.

**Fig 1.**
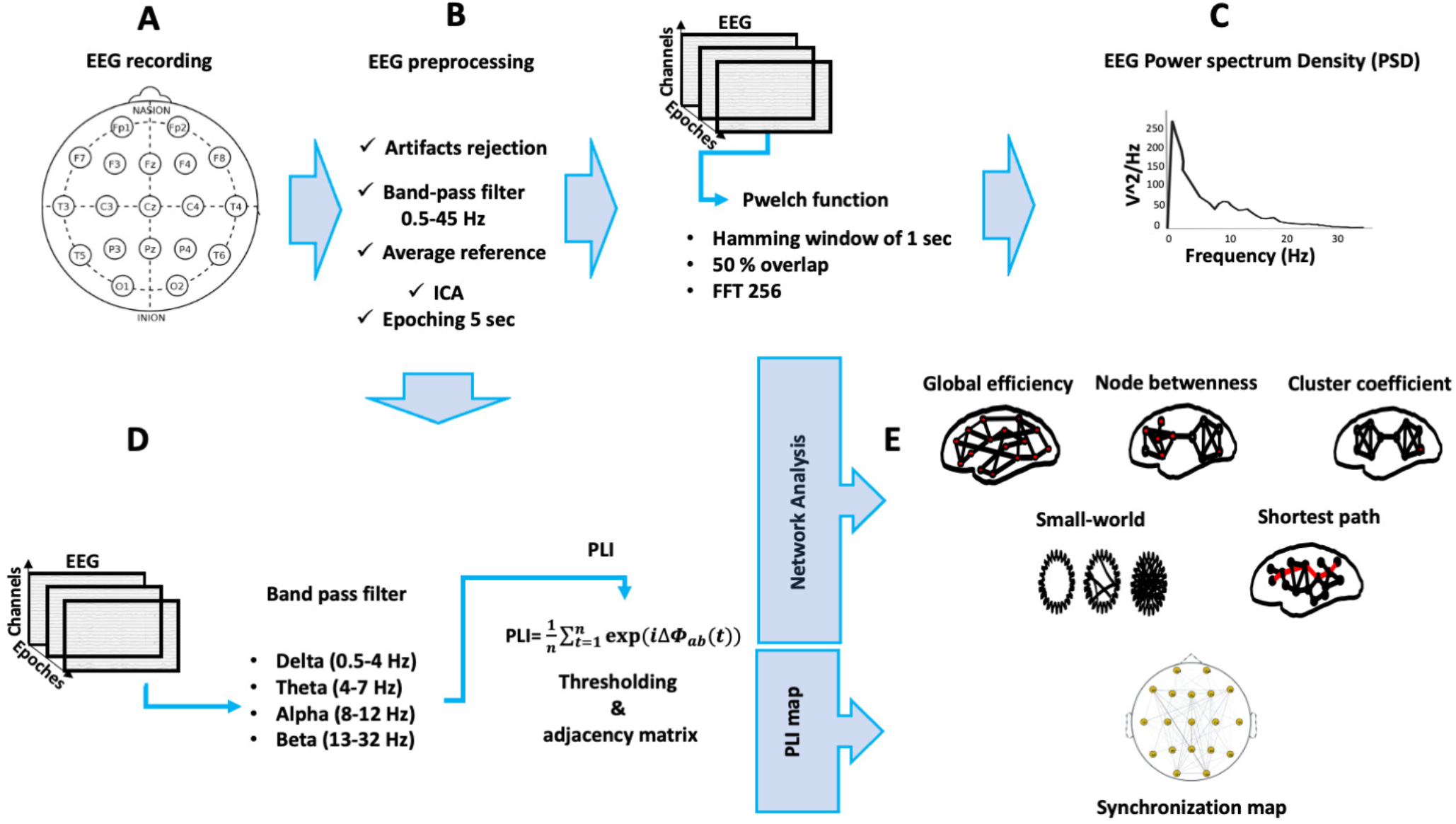
Flowchart of the proposed methodology. Panel (A) shows the international standardized 19 scalp EEG channels location. Panel (B) depicts the signal preprocessing pipeline used to clean the collected EEG time series. Panel (C) highlights the power spectral density processing step, where each single EEG channel was feed in the pwelch function. For each dataset, we extracted channel × epochs × frequency band. Panel (D) depicts the EEG based PLI network analysis. At this stage, each PLI matrix was thresholded and the adjacency matrix was computed to map the brain network graph. Panel (E) lists the indices used to test the performance of the network. In detail, we used global efficiency, node betweenness, cluster coefficient, small-worldness, and shortest path length like network indices.

### 3.3 Power Spectral Density Analysis

The artifact-free time series were submitted to a power spectrum analysis using the Welch periodogram method with Hanning tapered windows of 1s of duration and 50 % of window overlap. The Power spectral density (*μ*V^2^Hz) was estimated for a continuous range from 0.5 to 32 Hz in steps of 0.50 Hz. Our analysis was limited to frequencies lower than 32 Hz, why higher frequencies suggested very minimal contributions to those frequencies in the eyes-closed resting spectrum; in addition, we also tried to avoid excluding as much as possible electromyogenic activity. The power spectral density was computed using a multi-taper approach. Applying a multi-taper approach, the time series are sliced into segments and the average of their periodograms was found. For each time series x(n), the Welch’s modified periodogram averages a sequence of segments into overlapping sequences. In this paper, we use EEG segment equals to 5(s) length. This type of windowing is generally used to reduce spectral leakage and to smooth the power spectrum [26].

Let the overlap between the adjacent windows be represented as **K** samples, then the windowed r_*th*_ segment of data x(n) can be represented using Equation 1.

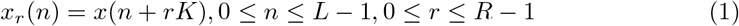

Where r has a length of 1280 samples.

The periodogram is calculated by computing the discrete Fourier transform and then squaring the magnitude of the Fourier transformed signal. The periodogram is then averaged, which reduces the variance of the individual power measurements. According to the Welch method, the power spectral density is expressed using Equation 2.

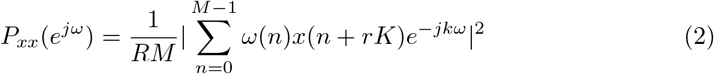

Where **n** is a number of slide windows and the length of each interval is M. R is the normalization value and can be calculated as depicted by Equation 3.

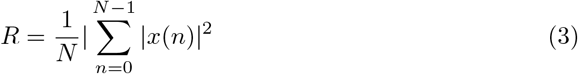

Since Welch method is the average of data sequences, the Welch’s power spectrum for each interval can be represented using Equation 4.

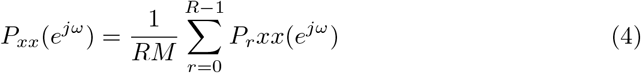

Where *P_xx_* is the periodogram computed on the EEG signal for each interval.

For this analysis, we use the “pwelch” function provided by MATLAB. To bypass the smears of energy in the spectral domain or the decaying tails, due to the convolution of the separate spectra of the signal, each r_*th*_ epoch of the sliced EEG signal was submitted to a Hamming window to reduce the spectral leakage and then processed by Fast Fourier transform to compute the frequency coefficient. The number of FFT points was assigned to 256. Finally, before applying further analysis, the EEG spectrum was transformed to decibel (dB) units (10log10(*X_n_*)) to standardize individual differences in EEG amplitude. To quantify an individual’s frequency, we further split the whole band 0.5 to 32Hz in the delta, theta, alpha, and beta bands with a maximum of 0.50 Hz within each narrowband under analysis. All PSD analyses were performed within subjects, resulting in channel × frequency × trial matrices of PSD values.

### 3.4 Graph Analysis

Graph theory offers a method to study the relation between network structure and function, regarding, for example, measures of efficiency, robustness, cost, and growth. A graph is a representation of a network, which is indicated by its nodes (‘vertices’) and connections (‘edges’). Graphs can be characterized by several parameters and principally by a clustering coefficient (CC) and shortest path length (SPL), node betweenness (NB), global efficiency (Ge), and small-world (SW). The small-world network architecture could be of primary importance for cortical dynamics because it represents a balance between local information processing and rapid sharing of this information with other regions. The common parameters to describe the graphs are the clustering coefficient (for segregation) and the path length (for the integration). The former is a measure of the local connectivity of the graph, whereas the latter is an index of overall connectivity. The small-world networks organization, instead, is focusing on an optimal balance between local specialization (segregation) and global integration. Therefore, we describe measures of functional integration and segregation and quantify the importance of individual brain regions, as well as to characterize local patterns and test the resilience of networks to PNES disorders. Graph analysis was implemented using the Brain Connectivity Toolbox [27].

### 3.5 Phase-locking index analysis

We apply PLI [28] to quantify the asymmetry in the distribution of the phase difference across all pairwise combinations of EEG sensors, resulting in a 19 × 19 (19 = number of recorded EEG channels). Finally, PLI between two electrodes was assessed getting the instantaneous phase from the analytical signal, obtained using the Hilbert transform as:

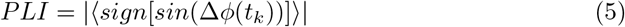

where Δ*ϕ*(*t_k_*) represents the phase difference and the **sign** function denote the average over time, **t**. Unlike other phase synchronization measures, such as phase coherence and the imaginary component of coherency, PLI is much less affected by the influence of common sources (volume conduction) and active reference electrodes [29]. Thus, under the assumption that the effects of brain volume conduction at the scalp level are instantaneous and no phase distortion is introduced, we compute the instantaneous phase of a time series *x*(*t*) as follows:

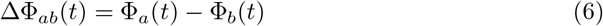

The equation 6 is denoted as the phase difference of signal **a** and signal **b**, where Φ_*a*_ is the phase of the signal calculated by Hilbert transformation. The PLI assesses the relative phase distribution’s asymmetry, which refers to the likelihood that the phase difference in the interval −*π* < Δ *ϕ* > 0 is different from the likelihood that it is in the interval 0*ϕ* < Δ *ϕ* < *π*. In this paper, we obtained the index of the time series with phase difference Δ *ϕ*(*tk*) (phase lag index) as follows:

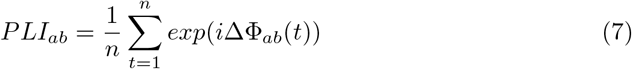

When ΔΦ_*ab*_(*t*) is a constant and the EEG time series is synchronized, PLI_*ab*_ = 1. If the time series is unsynchronized, then PLI_*ab*_ = 0. PLI values range between 0 and 1. The PLI value of zero indicates no coupling or coupling with a phase difference centered around (0*π*). The PLI value of 1 indicates phase synchronization in Δ*ϕ* different from (0*π*). The more consistent this nonzero phase locking is, the larger the PLI is.

In this paper, the complex network associated with each epoch_*i*_, was generated by measuring the functional connectivity (*PLI_i_*(*i,j*)) between all possible pairs of electrodes *i* and *j*. Considering *n* = 19 sensors, there are *n*(*n* – 1)/2 = 171 possible pairs of channels. The matrices of *PLI_i_*th were determined for every epoch (with *e* = 1,…, 240). The related (symmetric) connectivity matrix was generated by associating with each off-diagonal entry (i,j), the corresponding value of PLI(i,j). The functional connectivity analysis was performed as follows: for each patient, the original artifact-free EEG was subdivided in non-overlapping segments, here named (epochs); for each epoch, five 19 × 19 connectivity matrices have been derived; respectively in delta, theta, alpha and beta as well for the wide-band. To obtain a solid measure of functional connectivity, we used a 1 s length window to be within at least 4 oscillatory cycles. The lower edge was set to 0.5 Hz. Next, the network measures used in this paper were calculated on the basis of the 19 × 19 adjacency matrix. For each adjacency matrix, we calculated the graph G for each thresholded PLI matrix. Each graph is represented as *G* = (*N, W*), where *N* is equal to the number of EEG sensors and *W* = *w_ij_*; thus, *W* is the *N* × *N* symmetric matrix *w_ii_* = 0 and *w_ij_*. Finally, the PLI index, determined between electrode *i* and *j*.

#### 3.5.1 Network parameters

To perform network analysis based on PLI measure, the first step was to extract binary graphs, normalized to the interval 0–1, from the weighted connectivity matrices. However, it is noteworthy to mention that not all the weighted links in the initial connectivity matrices are significant, thus it is better to remove the non-significant branches and minimize the noise level. There are no exclusive methods for binarizing the connectivity matrices, but the binarization approach is the most used. A simple approach to binarizing a weighted connectivity matrix is to apply a threshold **th**, that is if a link has a weight higher than **th**, the corresponding entry value of the adjacency matrix is one, and zero otherwise. The common pitfall with the binarization approach is that several valid threshold values can be found to extract the important branches. To overcome this challenge, the threshold value was often included in a fixed range. However, there may be individual variations in the functional connectivity since some subjects might have higher average connectivity than others, thus the same threshold applied to all subjects may lead to different network densities. Many of the network topological properties can be biased by the binarized approach. Thus, to avoid these problems, one can study the network properties as a function of density instead of the threshold. It is common to use a threshold index such that the connectivity matrices have the same density values. One much-appreciated solution is to consider a range of density values and study the topological properties of the extracted networks. To obtain normalized network measures, the entry PLI values were normalized between 0 and 1, then were divided by the average obtained from a set of 101 random graphs (obtained by randomization of all actual matrices) with the same number of nodes and connections as the actual graphs.

For each subject under analysis, the adopted PLI-based network map consists of a fully connected graph with 171 nodes. The network size was stable over the epochs and the subjects. For every subject, the *PLI^e^* matrices are estimated, with *PLI^e^* = 1, …, 227. The entry PLI matrix, named *PLI^e^*, was binarized and the related adjacency matrix computed for every subject was computed. Now, let Ψ=*X_n_*∈R^(*r,c*)*N*^, be an EEG set holding N trials from the PLI dataset, recorded from a single subject, with **r** the number of sensors and **c** the number of samples of each trial. Here λ (*X*_*n*=1:19_→*Xn*′, *f_f_*) was a PLI measure between the phase of channel *X_n_* and *X_n′_* at frequency *f_f_*. We obtained a connectivity matrix Δ(*f_f_*). The value of **f** varies in the range from 0.5 to 32 Hz. Within the whole band we highlighted five sub bands here defined as delta, theta, alpha, beta and full 0.5 to 32 Hz. The PLI values range form 0 to 1, showing the degree of uncorrelated to fully correlated phase locking between pairs. At this stage, each trial is characterized as Δ(*f_f_*) ∈ *R^r,r,N,f^*, where Δ(*f_f_*) stands for PLI functional connectivity matrix. Next, statistical significance of PLI was determined to remove inconsistent or powerless connections. In this study, for each *N_i_* and *f_f_* we handled these data through three different threshold values (e.g 0.05, 0.15 and 0.25) that led us to get binarized PLI matrices. Finally, our Ψ was Ψ*X_n_* ∈R^*r,r,f,tN*^; where **t** is a number of threshold value (here equal to three). Numerically, for each subject (*X_n_*), we characterized a Ψ like # 19 × #19 × #3 × #5 × N, where # 19 are the number of EEG sensors, 3 are the threshold values and five are the frequency bands under investigation (e.g delta, theta, alpha, beta and full band). Each Δ(*f_f_*) matrices were averaged within bandwidth ff across **N** to reduce the final size of our Ψ to the final size of # 19 × # 19 × # 3 × # 5 for each subject.

### 3.6 Graph metrics

In the literature, we found a plethora of graph theory metrics, but not all of them are robust enough for studying the brain network functions. Here, we take into consideration several neurobiologically relevant network measures related to brain functions including information, segregation, and integration. Indeed, since EEG signals are interdependent, spurious connections may be generated by applying PLI measures to estimate connectivity from them. We tested the connectivity matrices over a range of different thresholds to overcome the issue of semi-arbitrary thresholding. We examined how the network density changed at various threshold values from 0.05 to 1 with step 0.1. The density of a network is characterized by the percentage of edges that remain after thresholding, which changes with the threshold level. The connection density from 0.05 to 1 with step 0.1 was tested as this was reported that 0.5 provides an optimal trade-off between reducing spurious connections and retaining true connections, providing biologically likely information about the brain functional networks. The graph analysis is explored in greater depth in section 4.2. Functional integration in the brain represents the ability to integrate information among distributed brain regions. Measures of integration or differentiation are developed to characterize the communication among brain regions. Measures of communication between cortical areas are commonly based on the concept of a path. The path is a sequence of distinct nodes and links, and in anatomical networks, represent potential routes of information flow between pairs of brain regions. We used a number of different graph measures including average shortest path length, global efficiency, cluster coefficients, small-worldness, and node betweenness.

#### 3.6.1 Averaged shortest path length

The path length is a measure of global integration within a network and represents the potential for functional integration between brain regions. The shortest path length between two nodes is the minimum number of edges between two nodes [30]. The average shortest path connecting any couple of nodes in the graph is the shortest path represented as follows:

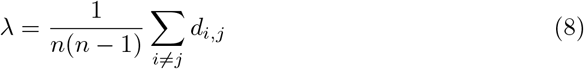

where *d_i,j_* stands for the shortest distance between nodes *i* and *j*. The averaged shortest path length is a graph property, which describes how well its elements are integrated/interconnected. A common drawback of the characteristic path length is that if any pair of nodes *i* and *j* are not connected through any path, the corresponding shortest path length value is *d_i,j_* = ∞. Here we refer to the averaged shortest path length as (SPL).

#### 3.6.2 Global efficiency

Differently from the characteristic path length, the global efficiency (*Ge*) is a measure of network performance based on its global topology. The global efficiency often increases by increasing network density (e.g, the number of connections), it is inversely related to the topological distance between nodes, and it measures the global information exchange across a network. Global efficiency is a graph index defined as the inverse of the average path length among all nodes [31]. Formally, it is calculated as follows:

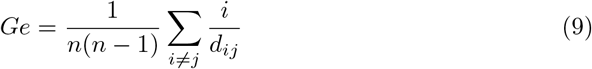

where **n** is the network size and *d_ij_* is the average distance between node **i** and **j** in the network. The global efficiency is a measure of the integrated information between distributed nets and stands for the overall network capacity to transfer information in parallel.

#### 3.6.3 Clustering Coefficient

Functional segregation represents the ability of processing that occurs within interconnected brain regions, and it is commonly applied to quantify the presence of clusters or modules within the network. Measures of segregation are based on the number of triangles (triples of fully interconnected nodes) in the network. Locally, the triangles of interconnected nodes around an individual node are known as the clustering coefficient. The clustering coefficient is equivalent to the fraction of the node’s neighbors that are also neighbors of each other [32]. It is a measure of local connectedness [33]. The mean clustering coefficient reflects the prevalence of clustering connectivity around individual nodes. The clustering coefficient *CC* quantifies the possibility that two neighbors nodes are neighbours. The average *CC* is defined as:

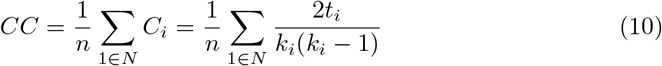

where *CC_i_* represents the cluster coefficient of the node **i** and *k_i_* is the degree of node **i**, and *t_i_* denotes the number of edges between pairs of nodes, **j** and *j′*, that are both connected to **i**. The number of the connections around a node **i** can be calculated as *k_i_*, (*k_i_* – 1)/2, where *k_i_* is the degree of node **i** [34].

#### 3.6.4 Small-worldness

The human brain consists of complex and specialized areas for sharing and integrating information [35]. The connections between cortical areas are not random, but are organized in a so-called small-worldness network topology. Integration and segregation properties combine functional specialization with higher-order processing like multi-sensory integration, cognition, and executive functions that require large-scale integration. The small-worldness reflects an optimal balance of functional integration and segregation [30]. The small-worldness network measure is defined as:

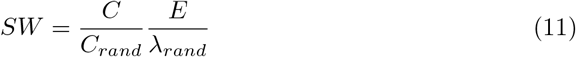

where **C**_*rand*_ and λ_*rand*_ are the clustering and path length coefficients of a random network with an equal number of nodes. The functional integration of brain regions (hubs) plays a crucial role in network interaction and communication across brain areas.

#### 3.6.5 Node betweenness

Node betweenness (NB) is a measure of the centrality of a node in a network and is calculated as the fraction of shortest paths between pairs of nodes that pass through the node of interest. Node betweenness is used to measure the influence of a node on the propagation of information through the network [36]. NB is defined as:

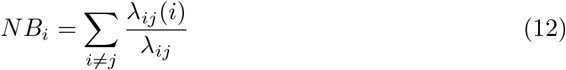

where λ_*i,j*_ is the number of shortest paths between nodes **i** and **j**, and λ_*i,j*_(*i*) is the number of shortest paths making use of the node **i**.

### 3.7 Statistical Analysis

To calculate the statistical significant differences between PNES and HC, we used a non-parametric cluster-based permutation test to address a multiple comparison problem. We investigated the relationship between brain areas and neural rhythms. First, we applied a Shapiro–Wilk test to grasp the distribution of our data. The null hypothesis is that the network data, in different frequency bands and at different thresholds, are independently distributed. Thus, we will accept the null hypothesis if the p-value is higher than the chosen significant *α* level. Additionally, our data were submitted to an independent sample, two-tailed permutation test based, using a family-wise alpha1 level of 0.05 and alpha2 of 0.001 to construct clusters of significant spatial t-value maps. Separately, we represent our results in box plot figures to highlight the differences between frequencies and brain area between PNES and HC. For the permutation test, we shuffled all epochs and splitted them into two datasets. Next, we constructed a cluster-level statistics and calculated a probability to estimate cluster-level p-values for each area and frequency. PSD data were the input of the cluster-level statistics. The output was a p-value for each cluster-level statistic. The cluster-level p-values were corrected and approximated using a permutation test. To test the impact of different threshold values in the network binarization methods, we applied a permutation test, we shuffled all the adjacency matrices to obtain a t-value map. We repeated this procedure using a simulation with 1000 iterations, and extracted the clusters from each permutation to compare the latter with the original dataset. Subsequently, we constructed a box plot of the cluster-level statistics with the calculated probability of cluster-level p-values for each EEG rhythm under analysis. In this case, we used three datasets of 3-dimension (eg. number of thresholds × number of parcellations × number of frequencies), as an input for the cluster-based permutation tests. We further conducted t-tests for the two datasets of network measures to obtain a t-value map of statistical significance in different frequencies and among the three brains under analysis. The analysis of the within-subject effects of PNES and HC was run with MATLAB 2021 and its associated toolboxes.

## 4 Results

In this section, we present the experimental results.

### 4.1 Relative PSD analysis

In the present resting-state EEG study, for each subject and each epoch, we have extracted a set of features. In particular, the relative delta, theta, alpha, and beta power bands were computed for each channel as the ratio between the sum of the original PSD (computed using the Welch method in the whole band 0.5–32Hz (total power)) and the preferred frequencies: delta (0.5–4 Hz), theta (4–7 Hz), alpha (8–12 Hz), and beta (13–32 Hz). Our power spectral density EEG analysis was also conducted within subjects. The background of the PSD decreased from slower to faster frequencies, according to an inverse power-law, with a **1/f**-like shape [37]. Notably, in the delta frequency band, the relative log normalized PSD values for PNES ranged in [0.3, 0.6].

In the beta frequency band, the relative PSD fluctuated in [0.025, 0.35], indicating that most energies focused at the lower frequency when the subjects were resting. Comparing the relative PSD values between PNES group and the HC (See Figure. 2, A), we found higher PSD in the delta band for PNES in sensors related to the frontal area. Conversely, in the theta band (See Fig. 2, B) the PSD results in a higher PSD pathway within the central to back area, whereas in alpha (See Figure. 2, C) the PSD it is generally higher for HC. In addition, our results underpin that compared to PNES, HC subjects show a statistically significant shift of spectral power from high to low in alpha frequencies in all the brain areas under investigation. Next, the PSD curves in the beta band (See Figure. 2, D) highlight a stable trend of oscillation between 0.1 and 0.3. To further grasp more insight into the power spectrum results, we performed a post-hoc analysis of PSD values in four frequency bands and three brain parcellations (Frontal, Central and Parieto-occipital areas) (See Figure 3). In this paper, we applied the following brain parcellation: we considered the sensors (FP1-FP2-F3-F4) as frontal; the sensors (C3-C4-Cz-Pz) as central; and the sensors (P3-P4-T3-T4-T5-T6-O1-O2-P3-P4) as parieto-occipital. The PNES group was found to have significantly higher PSD in the delta band in the frontal and central area (p < 0.05), in addition we found higher PSD in theta band respectively in frontal, central and parieto-occipital area (p < 0.01). In the theta band, post-hoc analysis showed that PNES group differed as compared to HC (**p**=0.0082, **p**=0.0476 and **p**=0.0076 respectively for frontal, central and parieto-occipital area). Differences was also present in the beta band (**p**=0.0255 and **p**=0.0308 and **p**=0.0043 respectively). In beta band we found higher values of PSD for HC compared to PNES. In the delta band, HC showed a lower PSD values as compared to HC (**p**=0.0262,**p**=0.342,**p**=0.0153). In alpha band the values of PSD for HC were higher than PNES with (**p**=0.0032, **p**=0.00443 and **p**=0.023 respectively) in all the brain parcellations.

**Fig 2.**
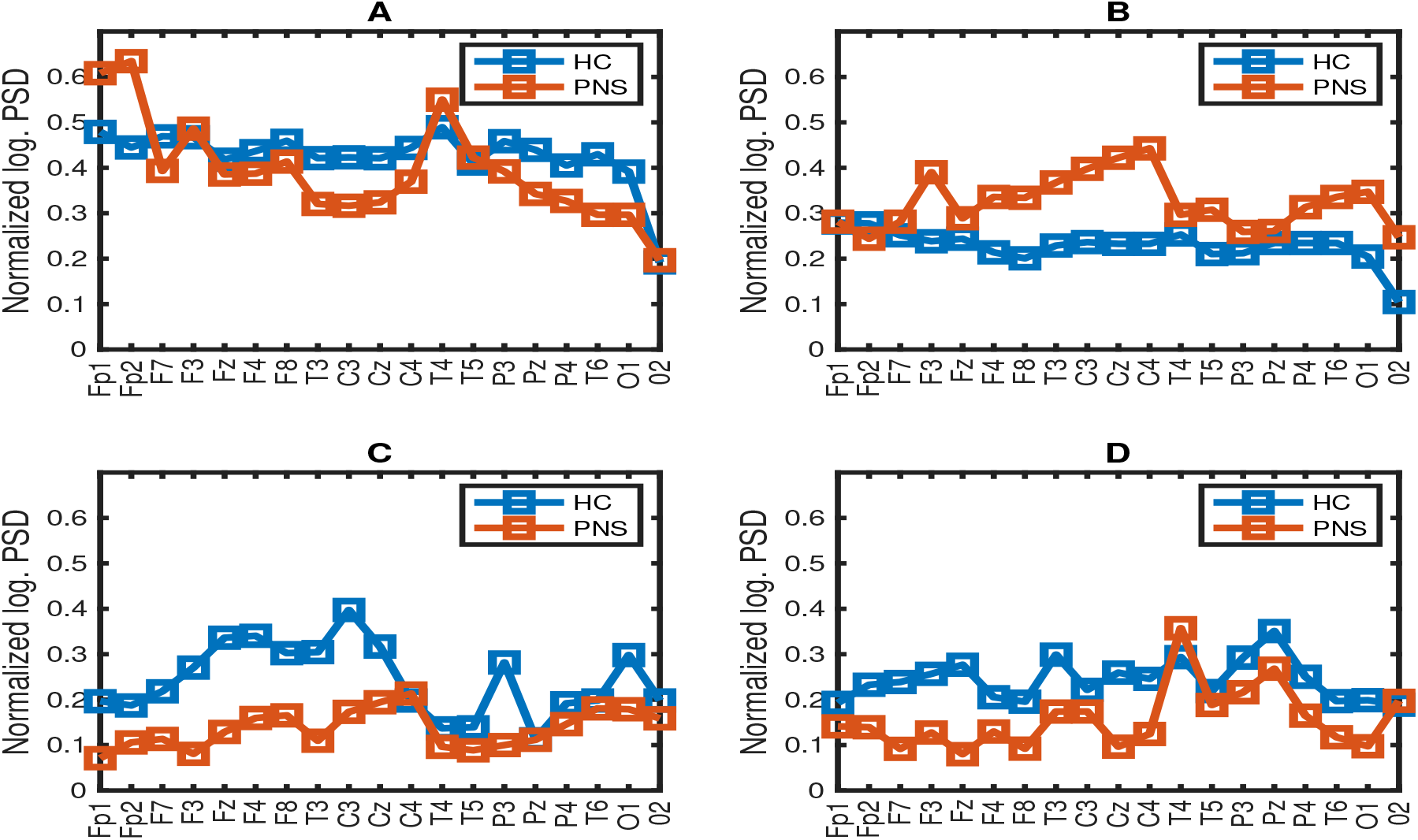
Grand-averaged of log normalized PSD of 19 EEG channels in the (A) delta, (B) theta, (C) alpha, and (D) beta frequency band for PNES group and the control group. Blue line denote HC subject and red line highlight the PSD of PNES

Compared to HC, the relative PSD values were increased for PNES in the delta band in the frontal and central areas. In contrast, PSD for alpha and beta bands were increased in the HC more than the PNES subjects in all the parcellations.

As illustrated in Figure. 3, frontal, central, parieto-occipital area displayed a significant PSD difference in the alpha and beta bands.

**Fig 3.**
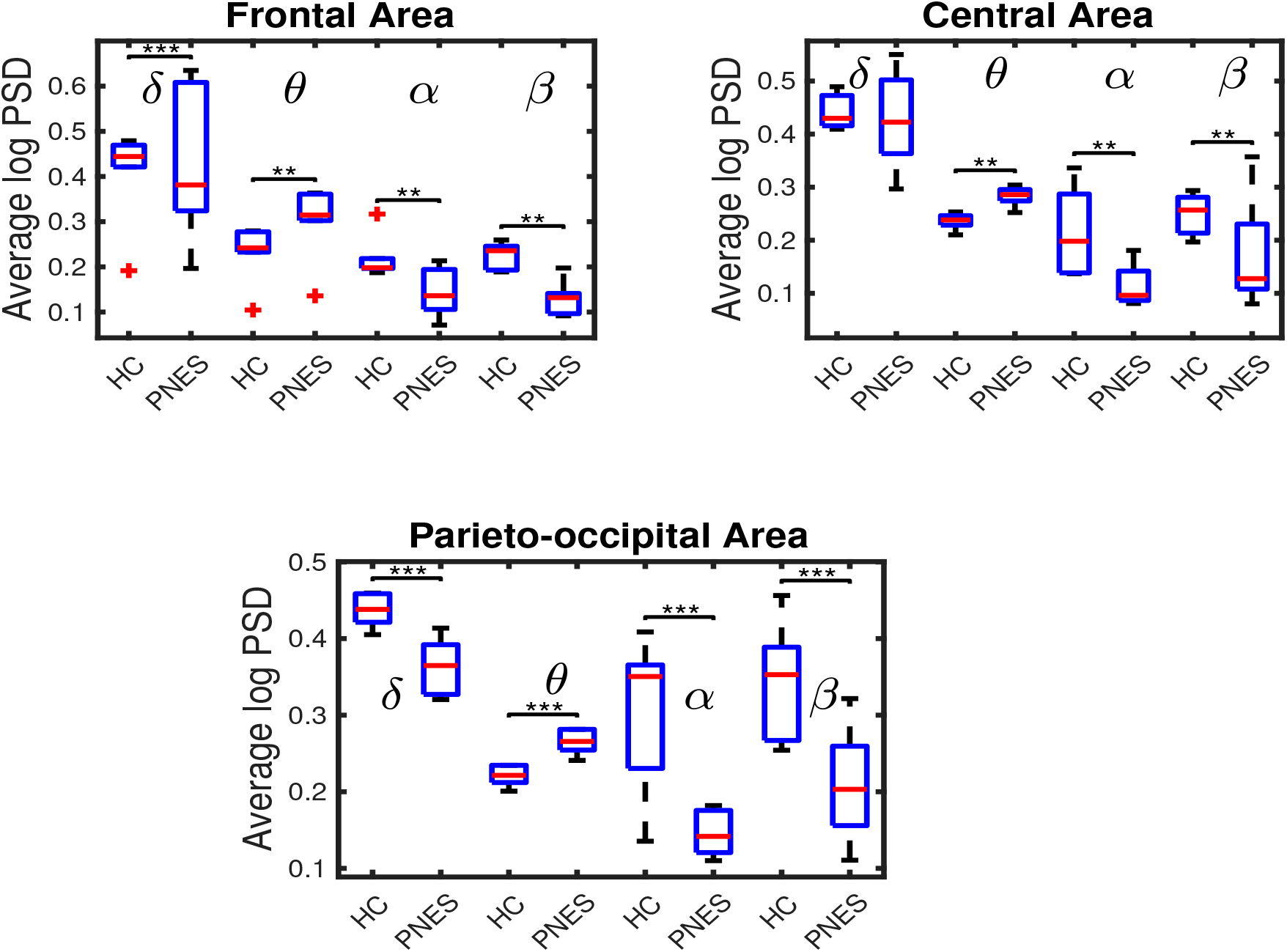
Relative normalized log PSD averaged over the 19 electrodes in four frequency bands for frontal, central, parieto-occipital areas between PNES and the control group. The horizontal line denotes significant difference between groups. ** **p** < 0.05 indicates that the p-value is statistically significant at *α* = 5%, and *** **p** < 0.001 indicates that the p-value is statistically significant at *α* = 1 %. The PSD measures was computed in PNES and control in the delta,theta,alpha and beta frequency bands. In delta frequency band significant differences were found between controls and PNES in frontal, central and parieto-occipital area. In the alpha frequency band the PNES had lower PSD value compared to controls. Differences were found in all the frequency band.

### 4.2 PLI analysis

To evaluate the functional brain network organization between the two groups, the networks were calculated for all frequency bands and for each epoch using PLI’s algorithm [28]. To grasp the differences in graph measures we performed a thresholding analysis at different values of **th**. A different threshold means a graph with a different edge density that lead to infer the degree of network disconnectedness. The disconnectedness of the graphs affects the quantitative values of many network metrics, thus choosing a good threshold might be a significant challenge in making a fair comparison between networks estimated in healthy volunteers and in PNES. All the graph metrics were initially estimated in the network from 0.05 to 1 with step 0.1 % of density.

We explored different thresholding values that force graphs to be connected even at sparse densities to address the issue of disconnectedness that can arise as a result of global thresholding. Furthermore, we attempted to dig statistically significant differences in graph indices at different thresholds in silico analysis of the real data. We computed a PLI analysis applying an iterative thresholding of the adjacency matrix (thresholding from 0.05 to 1 with step 0.01). Since the in silico computed curves of the network parameters are monotonous after the threshold of 0.25, we have chosen this value as the observation extreme (See. 4). To that end, we explored the effects of density on reliability by estimating all metrics in networks with different densities (see Figure 4).

**Fig 4.**
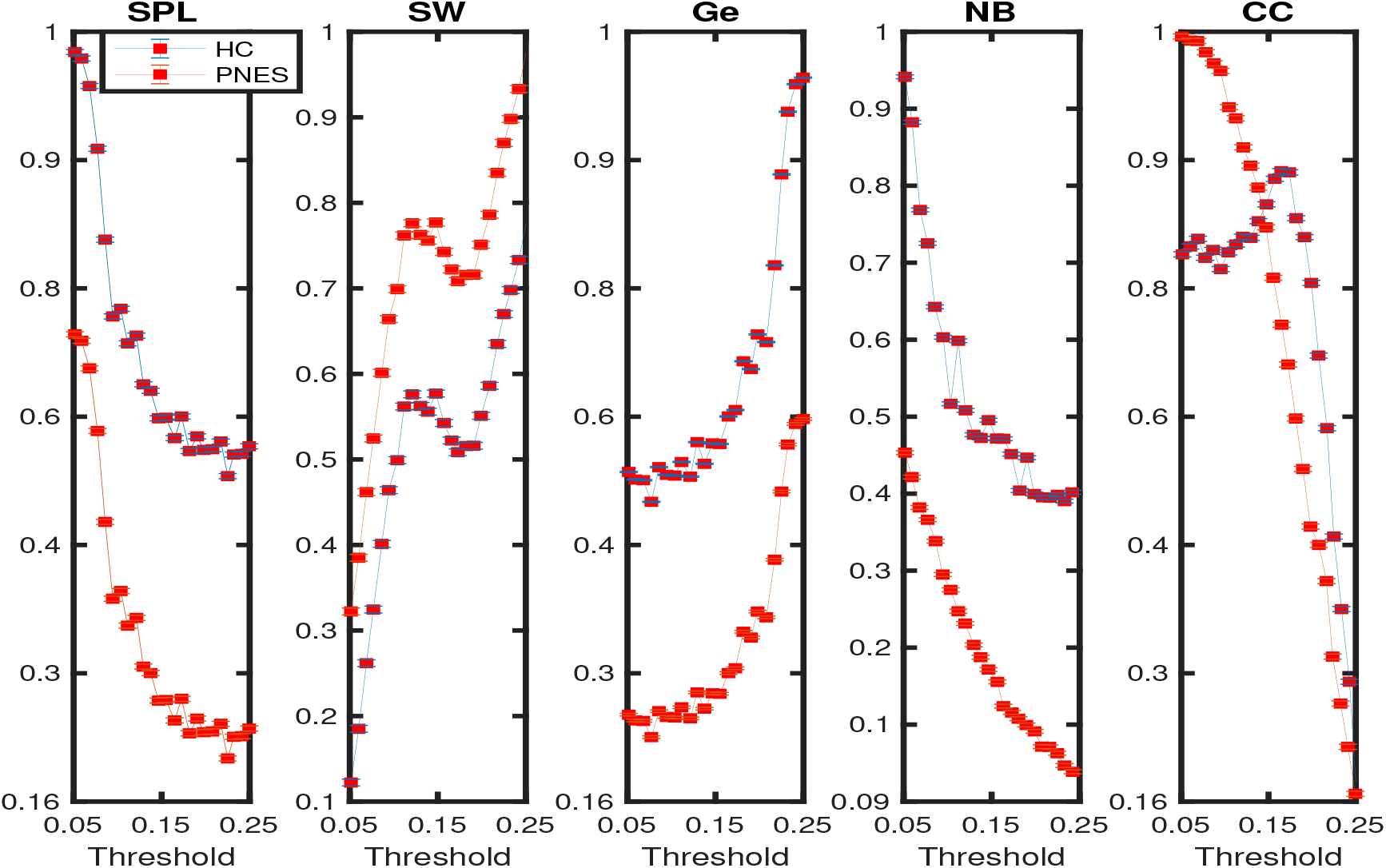
Network Density estimated by the PLI-based network analysis of the EEGs of the two group of subjects PNES and HC. Network Density was estimated by binarizing the connection matrix for different threshold values, as detailed in Section 4.2. This figure depict the mean and standard deviation results of clustering coefficient (CC), short path length (SPL), global efficiency (Ge), node betweenness (NB), small-world (SW) as the function of the thresholds 0.05, 0.15 and 0.25 at 0.1 step for the range 0.05 to 0.25. The analysis was computed on data in the whole frequency bands 0.5-32 Hz. The red colored circles, the blue colored squares are respectively for PNES and HC.

To explore how different values of the threshold could affect the network parameters, we explored three different points (here reported as 0.05, 0.15, and 0.25). In this way, we observe how the network parameters change as a function of the threshold and frequency. Additionally, to exclude the possibility that the observed associations between graph theory measures and thresholds of different frequency bands were biased by age of the subjects’ group as well as differences in brain activity, the EEGs was scrambled and subsequently processed as described (See Methods section). Noticeably, no correlations between graph theory measures and thresholds of different frequencies and age were found for the scrambled data. In Figure 5, 6 and 7 we summarize the results of different tests performed to find the differences between the network indices at different thresholds and between different frequencies between PNES and HC.

**Fig 5.**
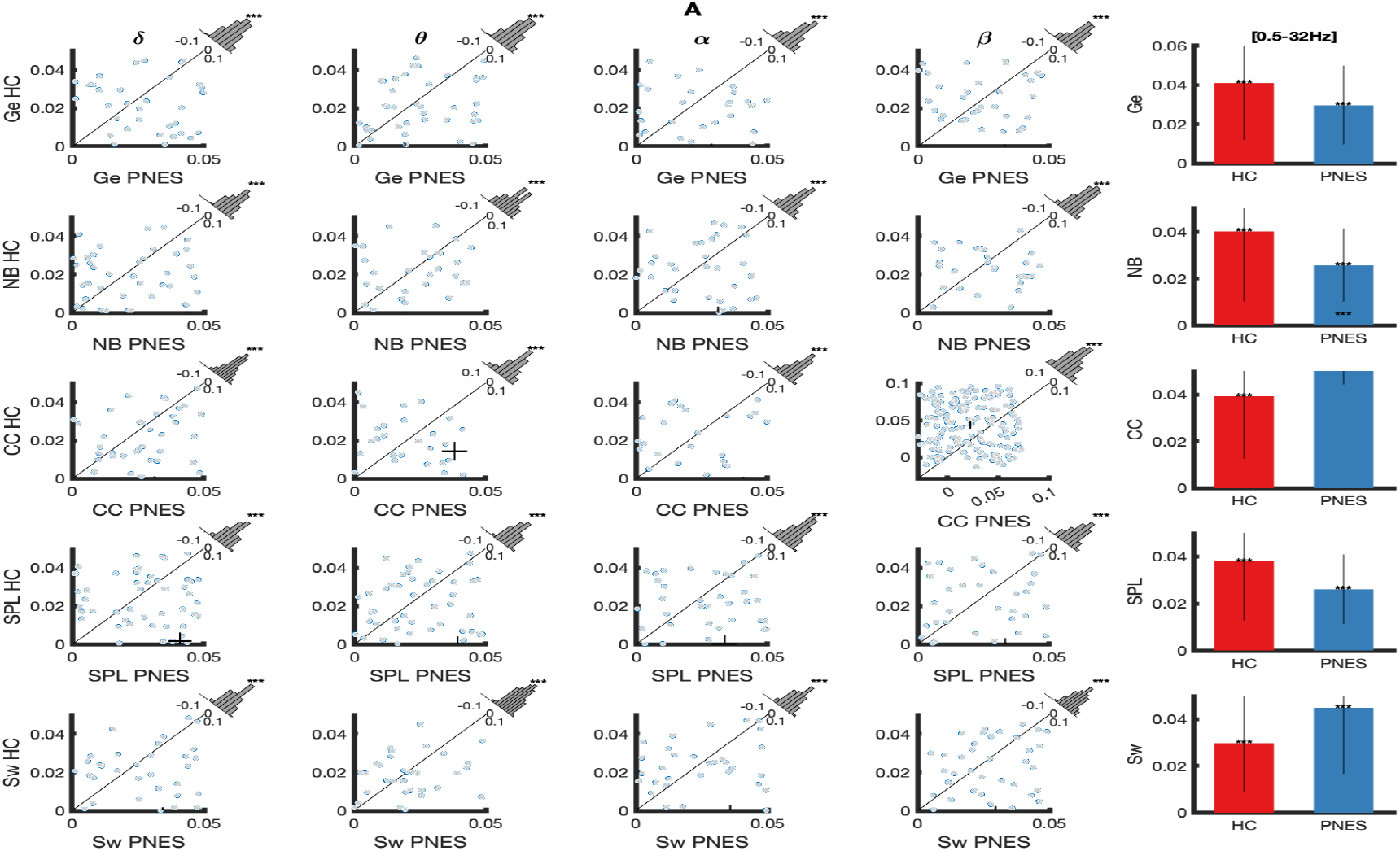
**Panel A:** The scatterplots depict point-like differences within specific band of frequency (e.g delta, theta, alpha and beta) for HC (x-axes) and PNES (y-axes). The point-like differences were computed as an euclidean distance between two network indices, PNES vs HC, at 0.05 of threshold. We used Euclidean distance to grasp differences in network measure distribution. Additionally, each histogram depict the data distribution and the **p**-value, and the significant samples are shown like a sample-specific t-tests at the Bonferroni-corrected level of 0.05. Last column of the panel show an histogram that compare the whole 0.5-32 Hz network indices value at 0.05 of threshold value between HC and PNES. Here the read and blue bins stand respectively for HC and PNES.

**Fig 6.**
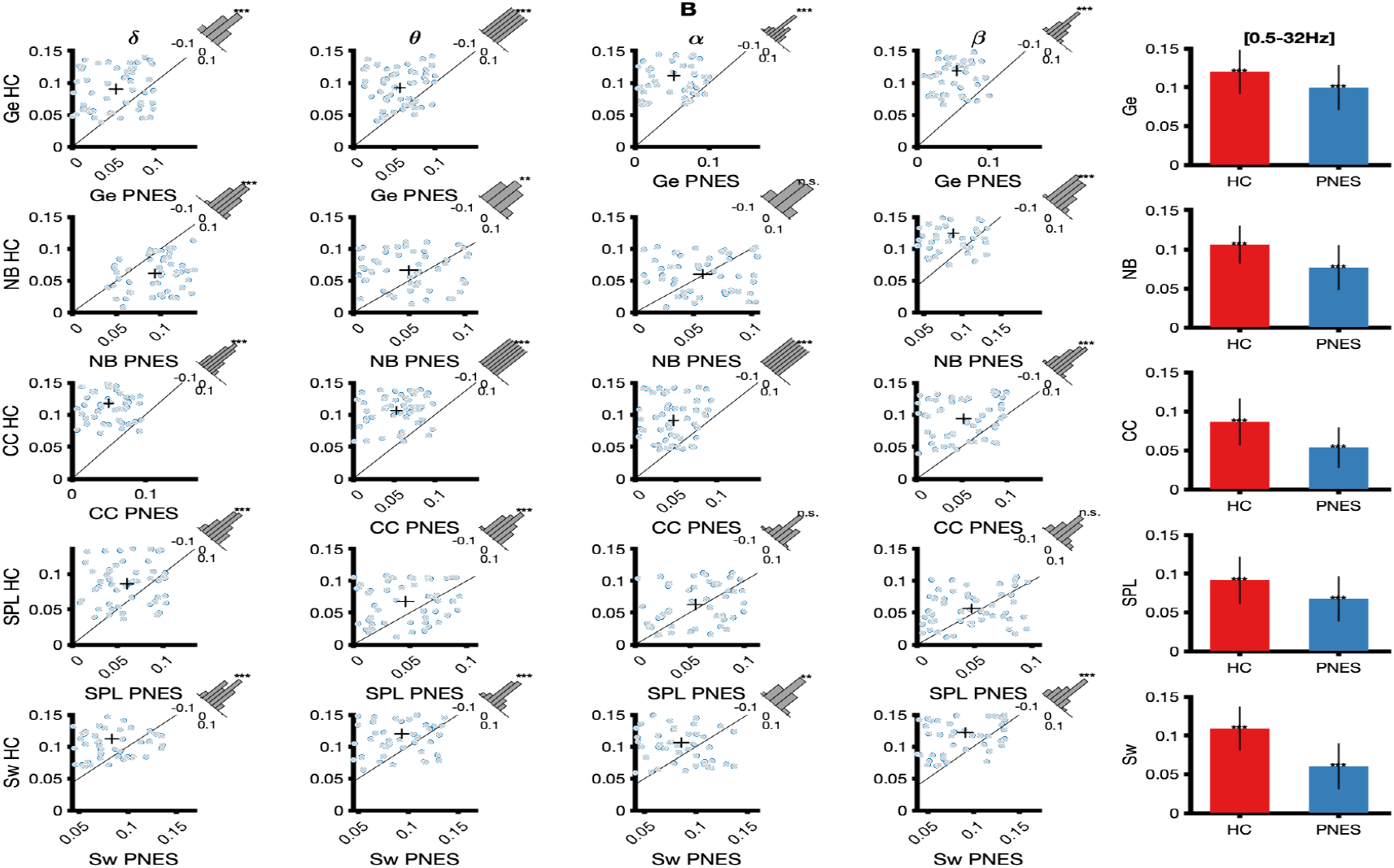
**Panel B:** Point-like distances of graph theory indices computed like an Euclidean distance between two class of subjects in each specific band of frequency at 0.015 of threshold. Here the raw of the panel rapresent a specific network indices, whereas the column a specific band of frequency. The scatterplots report the absolute values of distance measures among network indices and within specific band of frequency (e.g delta, theta, alpha and beta) for HC (x-axes) and PNES (y-axes). Additionally, for each scatter plot the linked histogram depict the data distribution and the related p-value, where the significant samples are shown as sample-specific t-tests at the Bonferroni-corrected level. The last column show an histogram that compare the whole 0.5-32 Hz network indices values. Here, the data were submitted to sample-specific t-tests.

**Fig 7.**
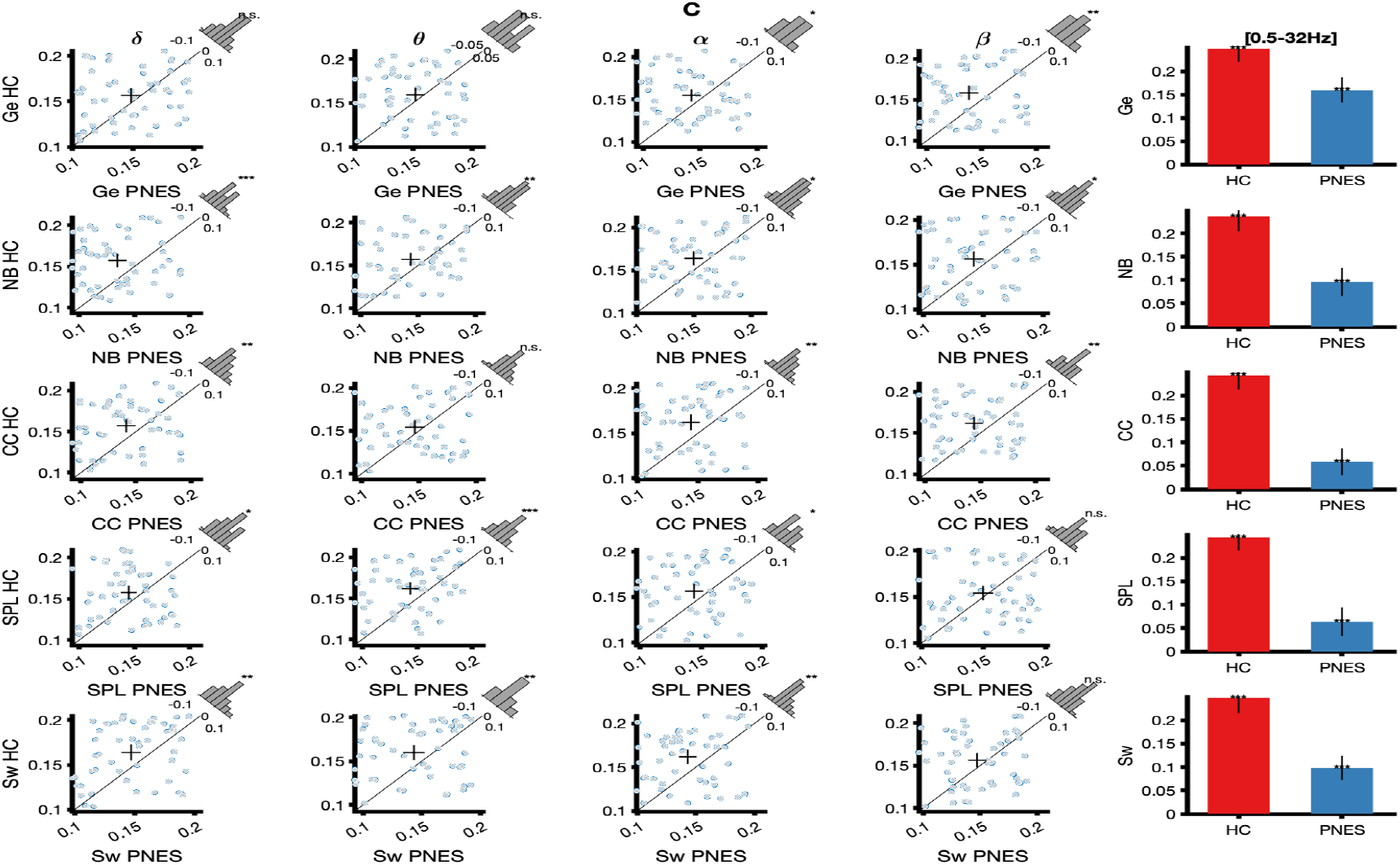
**Panel C:** Scatter plots of network indices difference (e.g Global efficiency, Node betwenness, Cluster coefficient, Small world and shortest path length). The point-like differences were computed as an Euclidean distance between normalized network values. Here the raw of the panel rapresent a specific network indices, whereas the column a specific band of frequency. The scatter plots report the absolute values of distance measures among network indices and within specific band of frequency (e.g delta, theta, alpha and beta) for HC (x-axes) and PNES (y-axes). Additionally, for each scatter plot the histogram depict network indices data distribution in that frequency and the p-value, where the significant samples are shown as sample-specific t-tests after Bonferroni-corrected

We calculated the Euclidean distance between specific network parameters to obtain a summary of all differences between the two classes. As reported in Figures from 5 to 7, the difference between the density of network coefficients calculated in delta, theta, alpha, and beta within three different threshold values provides information on the direction of the similarity/divergence in the network values. Differences between network parameters across threshold values in different bands between HC and PNES were computed using the permuted state vectors, and **p**-values were determined by comparing the observed statistics to the null (1000 permutations). To facilitate visual comparisons (See Figures 5, 6 and 7), the chi-square variance test was used by defining the category with the minimum distance as zero and the maximum depending on the calculated distance in each combination. For the comparisons, the level of significance dropped from 0.05 to 0.001 between the combinations. The Euclidean distance of the network indices (HC vs PNES) with statistically significant differences was depicted by each histogram. We found that all the network values were statistically significant (**p**=0.05) in a Monte Carlo random permutation test across all subjects within each frequency sub-band at 0.05 of threshold between HC and PNES. th(0.05)(Global efficiency; delta(**p**=0.045); theta (**p**=0.00032); alpha(**p**=0.032) and beta (**p**=0.0044)); (Shortest path length; delta(**p**=0.0133); theta (**p**=0.0024); alpha(**p**=0.013) and beta (**p**=0.033)); (Cluster coefficient; delta(**p**=0.023); theta (**p**=0.014); alpha(**p**=0.00023) and beta (**p**=0.0122)); (Small world; delta(**p**=0.0022); theta (**p**=0.032); alpha(**p**=0.0013) and beta (**p**=0.042)); (Node betweeness; delta(**p**=0.014); theta (**p**=0.023); alpha(**p**=0.0033) and beta (**p**=0.023)). Based on the aforedescribed framework we chose 0.05 as a threshold value for our analysis. Next, we used a method to compute the symmetry in the distribution of phase difference across all pairwise combinations of scalp sensors. The phase lag index can be considered as a measure of the synchronism of different EEG time series in the frequency domain. In this paper, PLI analysis was applied to all pairwise EEG channels for the PNES and the healthy control group. Based on the PLI method, the functional network can be reconstructed. When the entry of the PLI matrix (Figure. 1) is larger than the threshold, the entry is set to 0, otherwise 1. It is well known that many network measures are heavily dependent on network density. The approach is, therefore, to binarize the connectivity matrices in a way that produced networks having the same density [38]. We binarized the matrices using the threshold value of 0.05 (based on the aforedescribed test). The matrix *PLI_ij_* of 19×19 values was transformed into a binary matrix *A_ij_* (Adjacency matrix). Suppose the *PLI_ij_* value was less than the threshold of 0.05, the corresponding *a_ij_* of the binary matrix *A_ij_* is fixed to 1 (link), indicating the group difference of PLI between electrode *i* and *j* was significant; otherwise, it equals to 0 (no Link). Thus, the functional connectivity can be extracted from the PLI matrix, and the functional properties could be further quantified. Figures. 8, 9, and 10 show the topographic maps of the relative PLI for PNES and the HC group. The optimal threshold for each subject and frequency band was applied to the functional connectivity matrix to understand the global topology of large-scale brain synchronization in patients and controls.

**Fig 8.**
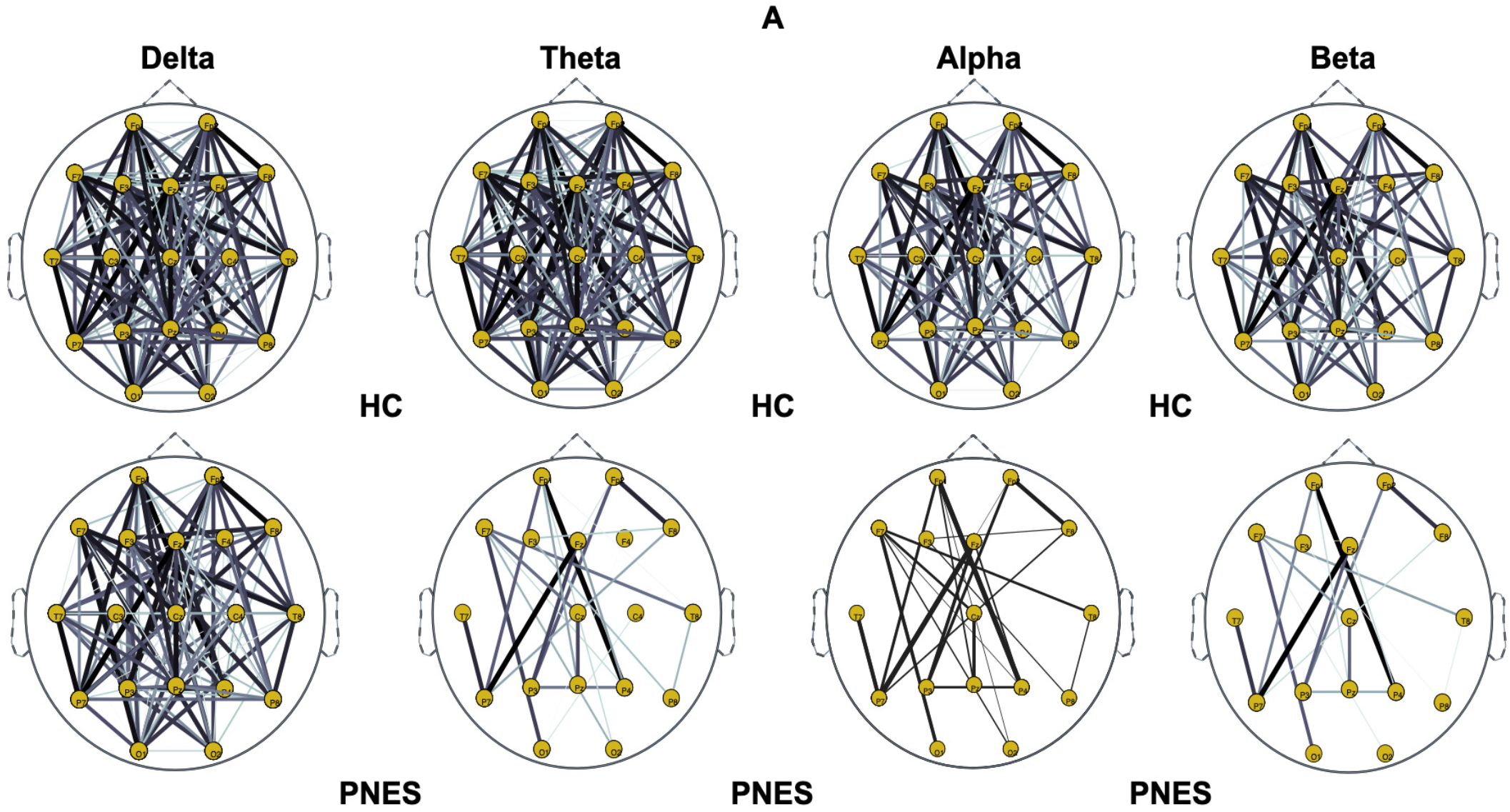
**Panel A:** Graphs representation of Phase Lag Index connectivity in Healthy subject and PNES in delta, theta, alpha and beta frequency bands. The channels layout in the network map reflects the location of the EEG montage in the acquisition system. The first couple of raw depicts the network maps of HC, the second one the PNES PLI maps tested at **th**=0.05. The line thickness highlights path with high value of PLI.

**Fig 9.**
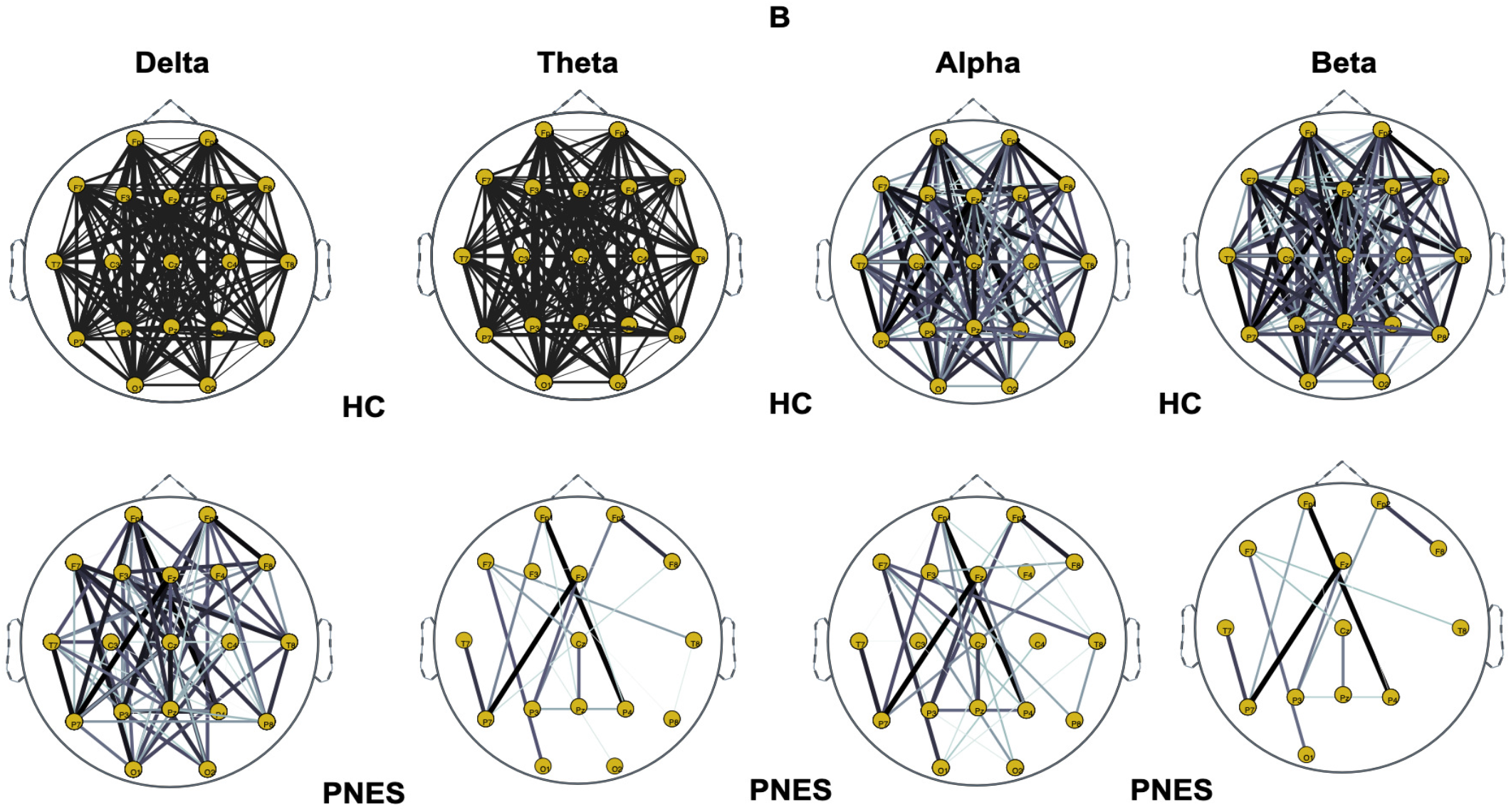
Panel B shows a Graphs representation of Phase Lag Index connectivity in Healthy subject and PNES in delta, theta, alpha and beta frequency bands. Here the PLI matrix was binarized using 0.15 as a threshold value. The first raw depict the network maps of HC, the second one the PNES network. Line thickness highlight path with high value of PLI.

**Fig 10.**
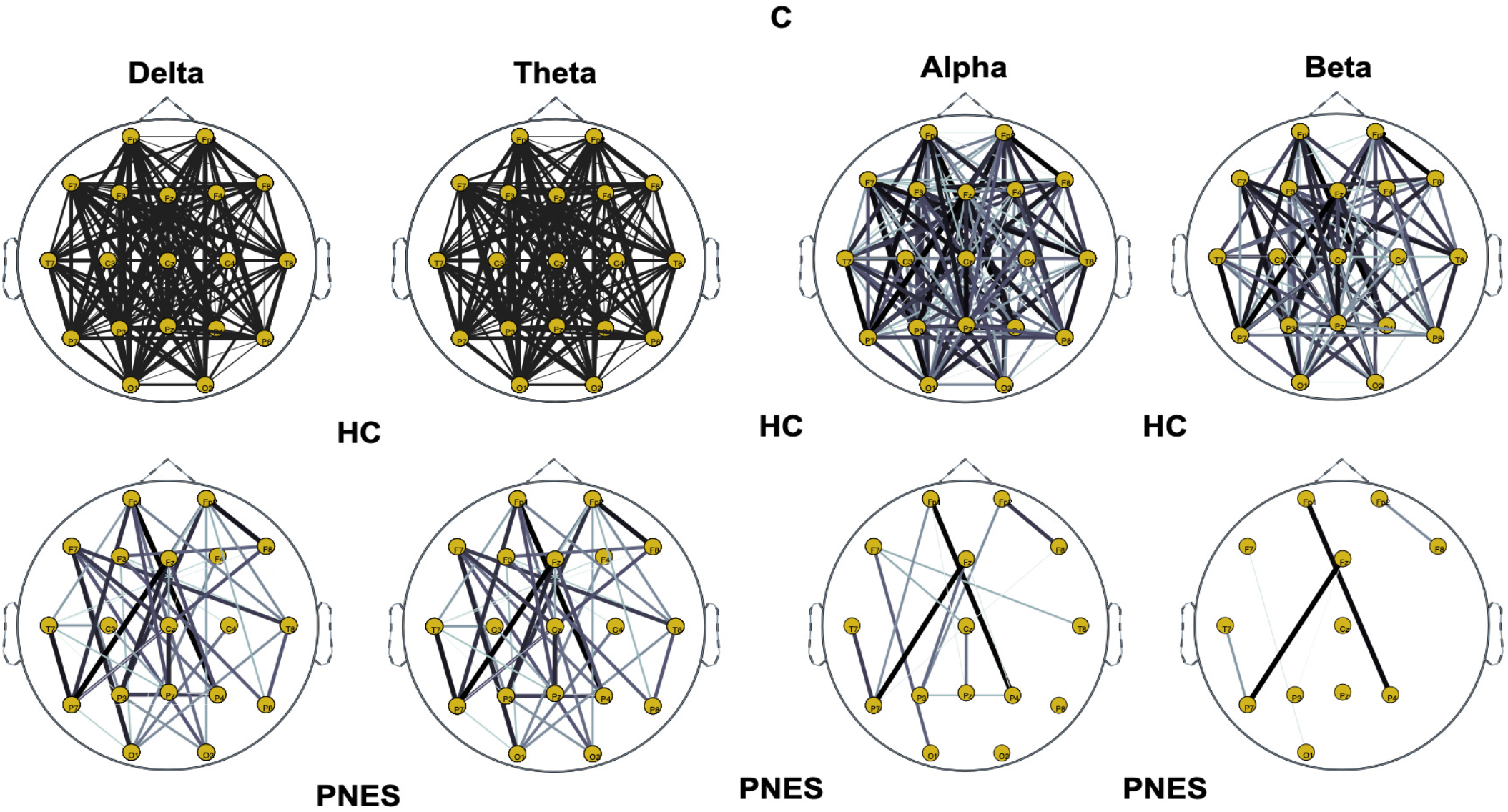
**Panel C:** Phase Lag Index connectivity in Healthy subject and PNES in delta, theta, alpha and beta frequency bands. The EEG channels layout in the map reflect the location of the EEG montage in the acquisition system. The first couple of raw depict the network maps of HC, the second one PNES tested using a **th**=0.25.

In the control group figures, 8, 9, and 10, there were more fully connected areas than in the PNES group. PNES group showed high values of synchronization in the delta and theta band, while scattered distributed connections in the frontal, temporo-parietal, and occipital areas in the alpha and beta bands were found.

HC group had a higher synchronization within frontal regions than PNES in delta and theta. We observed higher synchronization within the areas of the frontal and front-to–to-back regions for the HC group. In the beta band, less synchronization between the frontal, temporal, and parietal regions in the right hemisphere was observed for the PNES group. Lastly, in the alpha band, the synchronization in the left hemisphere was greater than the right hemisphere in the PNES group.

In Figure 11 we highlight the network coefficients measured in PNES and HC. We further performed a statistical test analysis based on two-tailed independent t-tests without assuming equal variance after giving F-test for the certification of group differences in the shortest path length, small worldness, global efficiency, node betweenness, and clustering coefficient at any threshold value accounted and among PNES and HC group. Our goal was to find important import differences that could lead us to grasp brain areas in PNES showing significant network impairments. Here, a significance level was taken as p < 0.05. Additionally, post hoc analysis showed that there was a significant influence of different frequency bands with bands and thresholds on node betweenness, cluster coefficient, and small worldness (all p-values corrected for double comparison using Bonferroni correction: p < 0.05).

**Fig 11.**
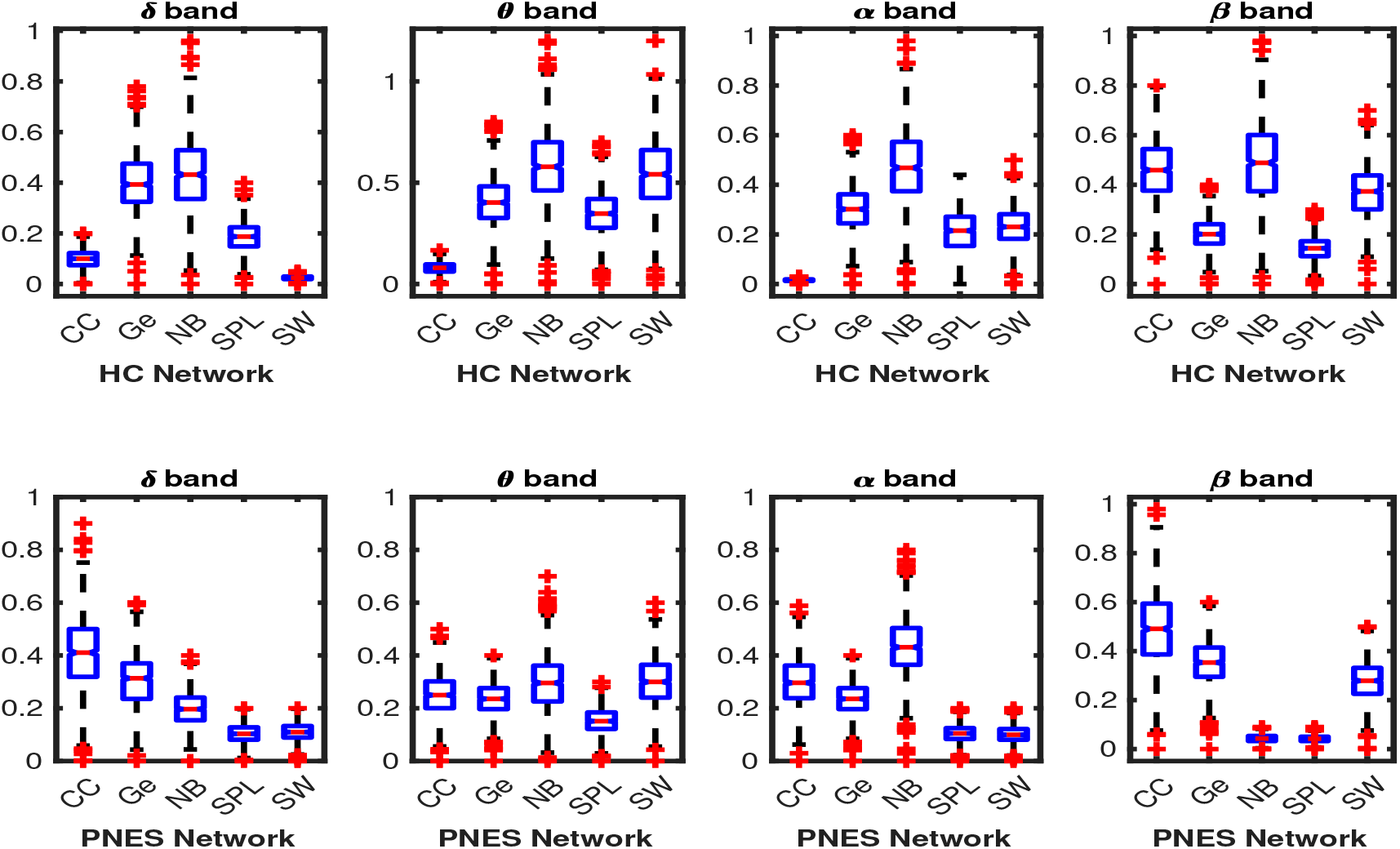
Box plots of network coefficients measured in PNES and HC. Here, we report normalized network coefficient values of: CC, *Ge, NB, SPL* and *SW* in delta, theta, alpha, and beta bands. The first raw highlights the network coefficients of the healthy control subjects. The horizontal mark within each box represents the median, the edges of the box represent the first and third quartile, the whiskers extend to the most extreme data points that are not considered outliers.

As reported in table 1, our multi-way statistical analysis depicted an overall significant difference in network indices in central and parieto-occipital areas. We have verified that beyond the threshold used to binarize the network, the areas of the brain in which the PNES exhibit a lower expression of complexity of network parameters were related to the central area and the posterior area. Furthermore, we found a higher statistical variation in the central area more than the parieto-occipital area. We found global efficiency differences in alpha (p=0.014) and beta (p=0.0011) with a central to posterior rate. Additionally, our results underpinned a Cluster coefficient statistical difference (**p**=0.0022) in alpha and (**p**=0.021) in beta for central and Parieto occipital areas. We found more small world differences in the central area than the occipital. The shortest path length is the network index with higher difference between brain areas, thus it could be the most discriminative.

**Table 1.**
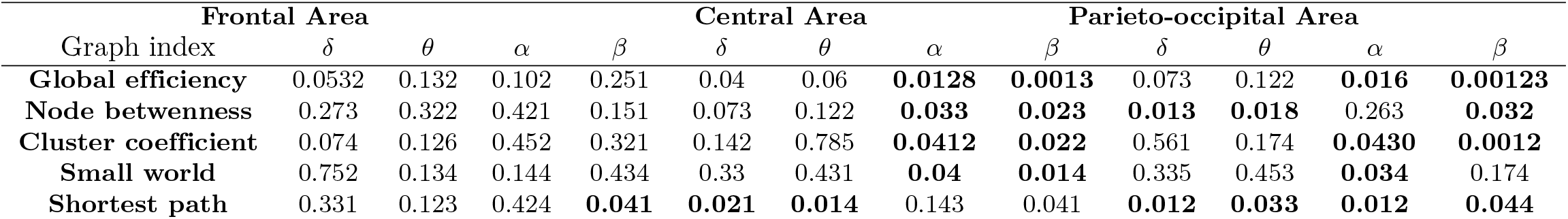
Results of our statistical analysis accross cluster of EEG sensors (See EEG layout parcelization in Section 4.1) in all of the frequencies band under analysis to grasp significant differences in network indices between the two classes (HC vs PNES). We evaluate Global efficiency, Node betwenness, Cluster coefficient, Small world, Shortest path characteristic. We performed a post-hoc analysis for multi-comparison. The **p** value (p<0.05) corresponds to significant difference in network index values for the testing conditions. See Section 3.7 for further information.

#### 4.2.1 Measures of integration and segregation

Measures of integration estimate the score of communication between distributed nodes. Specifically, the shortest path length (SPL) indicates that, on average, each node can reach other nodes with a path composed of only a few edges, while the global efficiency is an appropriate measure in the case of disconnected networks since the paths between disconnected nodes have infinite length, and thus zero efficiency. The global efficiency is the average inverse shortest path length, defined as one of the most elementary measures of the network’s integration but when defined only for the neighbours of a specific node, so this measure may be treated as an indicator of segregation that can share information between distributed regions. The measure of segregation defines the ability of specialized processing within a densely interconnected group of nodes in a specific network (clusters or modules). We also used a measure of segregation, the clustering coefficient, defined as the fraction of the node’s neighbors that are also neighbors of each other.

Segregation and integration are interdependent measures. Segregation decreases and integration increases as global pairing increases. When the global coupling is weak, there is high segregation and low integration, and the perturbed network nodes are disconnected and it operates in an independent way. Conversely, when the global coupling is strong, the integration is high and the segregation is low because the perturbed nodes are coupled.

We found significant differences in clustering coefficient (CC) in the delta, theta, and alpha bands. We found that CC was increased in the delta, theta, and alpha bands for the PNES group (Figure. 11. In addition, no significant differences are observed for Ge in all four-frequency bands (except Ge in the alpha). We found that Ge was relatively higher for HC group than PNES group in the whole band. As a measure of integration, the shortest path length shades light over the efficiency of global information communication. The shortest path length is a unique sequence of edges that connects two nodes, and its length is given by the number of steps or the sum of the edge with the shortest path length. We found a decrease of the shortest path length in delta, theta, alpha, and beta bands for the PNES group compared to HC group. In the beta band, the SPL value is slightly lower in the PNES group than HC group (Figure. 11). The human brain can be described as a small-world network structured around a large number of spatially distributed network communities with clustered connectivity, in which the local computations are highly segregated. In small-world architecture, integrated and segregated information are promoted by network hubs that arrange efficient communication and information processing.

#### 4.2.2 Measures of Centrality

The centrality is a measure of the relative importance of a node connecting a network in the brain and hence it is important for the functional integration of the brain. The different measures of centrality exist, including degree, closeness centrality, eigenvector centrality, and betweenness centrality. Nodes with high centrality are called hubs. Our results highlighted a decrease of Node betweenness in PNES in delta, theta, alpha, and beta bands more than HC group (Figure. 11). Our findings highlighted a loss of network performance.

## 5 Discussion

Different cortical brain regions are toned to oscillate at an individual rate [39]. These oscillations reflect local, intrinsic physiological mechanisms related to the fine-tuning of corticothalamic circuits within which each cortical region tended to resonate at approximately the same frequency. Rhythmic patterns of neural activities are believed to play an important functional role in local processing and communication between different neuronal systems. Thus, the study of brain frequencies in PNES does not only have theoretical relevance but also clinical implications. In addition, we used PSD to find specific differences in modulation in frequency tuning of brain areas among PNES subjects and healthy control [40].

We further investigated the functional connectivity in brain networks of PNES patients and healthy controls. The functional network represents the temporal lag of phase in the brain network communication, and the network measures are tools used to investigate the efficiency of communication between brain regions. The human brain can be described as a small-world network, that is structured around a large number of spatially distributed network communities with clustered connectivity, in which the local computations are likely to be highly segregated. Integration of segregated information in small-world is promoted by network hubs, which connect network communities and ensure efficient communication and information integration. Our results are discussed in the next subsections.

### 5.1 PSD measures

Figure 2 shows that the PNES subjects and controls had a similar pattern of the power spectrum. However, we observe that the relative PSD of PNES patients are significantly increased in the theta frequency band (See Figure 2 B) while markedly decreased in the alpha frequency band (See Figure 2 C). In Figure 3 we check for a spatial pattern of PSD to infer brain areas with strong differences across PNES and control subjects. Particularly, we found that in central and parieto-occipital areas the PSD of healthy subjects was higher than that of PNES subjects. We hypothesize that these results in HC may be related to a wealth in information processing and network connection and that the neural model of the healthy subject engages a greater number of processing and exchange of information in long as well in short-range. On the other hand, an increase of PSD in delta band in frontal and central areas for PNES may reflect a lack of influence of subcortical structures on cortical activity [41]. Frontal and central areas are implicated in the control of sensory feedback and attention. The hypothesis that in subjects with PNES there is a greater increase in power in these areas, leads us to think that there could be a greater alert and pre-activation of the cortical circuit implicated with sensory and movement control. Thus, the pre-activation or hyper-activation in this cortical circuit could leave this to fire and trigger a long front-to-back route when activated. Our findings, supported by other studies [18, 19], highlighted that PNES network alterations are related to high-frequency oscillations. Additionally, we found alpha rhythm alterations in PNES in different brain areas. Therefore, alpha alterations could reflect pathological resting-state dynamics in which thalamus, midline, frontal and parietal cortices play an important role in accountability to seizures [42]. The alpha rhythm is thought to arise through cortico-thalamic interactions, and to possibly reflect top-down processes that subtend a vast number of cognitive operations, in particular attention, working memory and sensory system control [43]. Arıkan et al. [19] found that PNES correlates with high-frequency oscillations on central and parietal areas, whereas Meppelink et al. [18] found a decrease in PSD in the beta band.

### 5.2 PLI Measures

In Figures 5, 6 and 7 and in Figures 8, 9 and 10 we summarize our analysis in network measures at different thresholds. We tested the network density of shortest path length, small world, global efficiency, node betweennes, and cluster coefficient in the range 0.05 to 1 at 0.01 steps. Our results highlight, (See Figure 4), how these curves assume a shape of a monotonic function. Additionally, we found that 0.25 is the knee point after which the curves assume this characteristic. We also observe how the curve maintains a certain degree of separation for each threshold against which they have been tested. In Figures 5, 6 and 7 we performed an Euclidean based difference analysis between coefficients data to grasp how the index differ between two groups. In table 1 we present our pos-hoc analysis to highlight differences at different thresholds in delta, theta, alpha, and beta for the graph measures are used. These results may suggest frequency-specific network organization and imply various functional roles for different frequency bands. Additionally, we highlight that in delta, alpha, and beta all the network metrics tested at 0.15 and 0.25 of density were not enough significantly different. Furthermore, we observe that 0.05 is a consistent and reliable value of threshold to investigate the resting-state graph metrics of PNES with lower bias.

Recently, the scalp EEG-based studies revealed that PNES might lack a relatively long linkage in the brain network topology, indicating the impairment of information transferring and processing [8]. As shown in Figures 8, 9 and 10, the two network topologies differ in the bilateral hemisphere with a marked difference in alpha and beta band. Decreased functional connectivity in the alpha band could be associated with a decreased attitude to generate seizure oscillations in EEG rhythm, thus uncontrolled movements of the PNES subject could be triggered by altered neurobiological substrates. This hypothesis is corroborated by the finding that PNES patients exhibited a more path-like topology with a decreased long-range synchronization in the alpha and beta band. Moreover, the network properties are effective to capture the information of spatial differentiation in the networks. The network topology differences in Figures 8, 9, and 10 shows that subjects with PNES have fewer and weak links between the frontal and the temporal/occipital areas than subjects in the healthy group, which is consistent with the structural abnormality difference between the two group found in [44, 45]. The working mechanisms of PNES are further revealed in Figures. 8, 9, and 10, which shows that the PLI analysis captures the differentiating spatial network information in the brain network. These results corroborate the hypothesis that PNES is a phenomenon of neuronal disconnection and that in high frequency, integration and differentiation of the network are lost.

### 5.3 Measures of Segregation and Integration

In this paper we used CC, Ge, and SPL to evaluate segregation and integration. The boxplot in Figure. 11 shows how the PNES subjects have increased clustering coefficients and decreased shortest path length as compared to the normal subjects; this could be a predictor of impaired local information processing and loss of global integration in patients [46]. In PNES the higher clustering values are a feature of regular local networks with symmetrical structures and highly connected with their closest neighbours. Instead, short path length describes the number of edges among vertices, and we utilize it as a feature known for random networks. PNES has been shown to correlate with functional and structural brain network alterations using MRI-related studies [12]. We also found a decreased global efficiency that could reveal that PNES might lack a relatively long linkage in the brain network topology, indicating impairment of information transferring and processing. In practice, PNES patients can be easily discerned from normal subjects based on their abnormal behavior. The CC, Ge, and SPL are measurements determined by spatial topology where the complete information of a network is significantly more complex than the adopted statistical measurements. Thus, the essential spatial information of a network would be meaningful to further improve the ability to differentiate the PNES subjects. In detail, we found a decreased averaged shortest path length (SPL) in PNES for all the EEG sub-bands under consideration and Ge in the delta, theta and alpha bands, indicating the network of PNES subjects like a model highly segregated and unable to integrate information over long distances. The general hypothesis is that to process relevant information to a momentary challenge, the neural responses are not only reflected by a change of neural activity in certain regions of the brain but also by a global reorganization of connectivity patterns. We highlight a loss of network synchronization (See. Figure 11) in the rest of the EEG of PNES as a spatial pattern related to a change of neural activity in certain regions involved in a momentary challenge and a global reorganization of connectivity patterns during the information processing. We argue that functional measures of segregation and integration can improve the knowledge on the rest behaviours of the whole-brain of PNES subjects. We can evaluate the ability of the brain to encode information using CC, Ge and SPL measures as well to characterize the effective integration of distributed information across the whole brain 11. The measures of connectivity calculated as the strength of the edges between the nodes highlight loss of functional integration between brain regions due to the loss of long-range linking. In addition, our results are in agreement with other studies [5, 7, 10]. Different from Xue et al. [8], we report cluster coefficient values higher in PNES than HC. However, we also observe that PNES subjects had higher CC than HC.

### 5.4 Measures of Centrality

We used centrality to characterize the relevance of individual nodes in the network using Node Betweenness. Our finding revealed that HC had a higher level of NB than PNES subjects, except for the beta band. Furthermore, our analyses suggest that impaired information processing involves connectivity across multiple cortical areas in the beta bands. Moreover, for PNES, in the beta band, fewer brain connections were found in the right frontocentral cortical areas. This is coherent with previous literature evidence, pointing to the pathogenesis of PNES disorders resulting from the contribution of the right hemisphere [20, 47, 48].

### 5.5 Small-Worldness

In order to further understand if clustering coefficient or path length are contributing equally to the observed small-worldness differences between group we performed a small-world analysis. The small-world networks have a high clustering and a short path length like random networks. Given that the small-world model supports both specialized and integrated information processing in the brain, we attempt to describe the effect of the impairment on the topological property of small-worldness characteristics derived from EEG connectivity data. Furthermore, the PNES group present a significant network differentiation in theta, alpha, and beta bands, with fewer small-worldness characteristics. As reported in Ding et.al (2013), the PNES small-worldness had altered functional and structural connectivity networks, which is related to a more regular organization in large-scale brain networks. Additionally, Ding et.al (2013) found alteration in region involving attention, sensorimotor, subcortical networks [12]. In other studies, the loss of integration and segregation in PNES was supposed to be related to emotion, executive control, and motor function impairments [10].

## 6 Conclusion

In this study, the proposed methodology can reveal subtle changes in the graph properties of PNES at different EEG frequency bands. We found that PNES patients present an increased spectral power density in the delta and theta bands in the frontal and central areas and an overall decreased spectral power in the other frequency bands under analysis. Additionally, functional connectivity networks exhibited altered nodal characteristics in global efficiency, node betweenness, and path length. We also found a decreased coupling strength of functional connectivity in PNES. The synchronized oscillatory maps showed high sensitivity to differentiate PNES patients from healthy controls. It supports the hypothesis that the PNES reflects an irregular brain architecture with a loss of functional connectivity and a disturbance of the right back-to-front or vice-versa pattern of information flow in brain areas related to cognitive operations, attention, working memory, and movement regulation. Additionally, as reported in table 1, we found that the shortest path length and node betweenness are the most significant/discriminative indices with differences within high frequency in the central and posterior areas. Differences in network parameters could highlight impairing in segregation and integration as well as in information processing during rest activity at the time of Video EEG monitoring. These alterations could also be related to the psychological condition, stress induced by the transient EEG monitoring event, as well as the alertness of the subject waiting for a particular/apparent trigger. From this perspective, PSD and graph measures might be effectively used to discriminate PNES in scalp EEG time series. As future work, we will attempt to increase the cohort of PNES subjects as well to collect EEG data with an high density acquisition system to raise the statistical precision of our analysis. It could also be interesting to investigate whether the abnormal organization found in this study can be further confirmed in a large cohort of PNES subjects using other methods such as weighted PLI or directed PLI. Next, we plan to apply this method to data acquired during transcranial magnetic stimulation (TMS) [49] to study how the brain networks of PNES changes under external perturbations.

## Author Contribution

**SG, SN** and **UA** Data Curation. **GV** Conceptualization, Resources, Software, Investigation, Formal analysis, Writing & Editing Original Draft. **GV, WB, BB, SN, MLG** Revision and editing. **GV, WB, BB, AH, NM, CI, FCM, MG, KD** Revision final version. All authors read and approved the final manuscript.

